# WasteFams: A database of protein families from global wastewater microbiomes

**DOI:** 10.64898/2026.05.08.723720

**Authors:** Alexandros Galaras, Iro N Chasapi, Eleni Aplakidou, Maria N. Chasapi, Efthimia Lamari, Sophia Diplari, Ilias Georgakopoulos-Soares, Evangelos Karatzas, Fotis A. Baltoumas, Nikos C. Kyrpides, Georgios A. Pavlopoulos

## Abstract

Wastewater surveillance has emerged as a critical tool for global epidemiology, yet the functional diversity of wastewater microbiomes remains poorly characterized at the protein level. Here, we present WasteFams, the first comprehensive database dedicated to the systematic exploration of protein families in wastewater metagenomic and metatranscriptomic studies worldwide. Integrating data from 580 metagenomes, 132 metatranscriptomes, and 1,709 reference genomes, WasteFams catalogs 3,887 non-redundant protein families (containing ⪰100 members) derived from over 105 million predicted proteins. Each protein family is enriched with multi-layered annotations, including AlphaFold3 structural predictions, taxonomic classifications, and biome-specific metadata. To further expand their functional annotation, we integrated deep genomic context analysis to link protein families to Mobile Genetic Elements (MGEs), Biosynthetic Gene Clusters (BGCs), Antibiotic Resistance Genes (ARGs), and CRISPR elements. Accessible through the EnvoFams portal, WasteFams provides a user-friendly interface featuring advanced search capabilities, sequence and structural similarity tools, and interactive visualization modules. As global initiatives increasingly leverage wastewater for public health and environmental insights, WasteFams can serve as a critical resource for discovering novel microbial functions, monitoring resistance mechanisms, and exploring the biotechnological potential of secondary metabolites within wastewater-engineered ecosystems.

## INTRODUCTION

Wastewater surveillance has emerged as a powerful strategy for monitoring public health and human activity at the population level, with the COVID-19 pandemic highlighting its value for global epidemiology (1). Because it is cost-effective and non-invasive, wastewater monitoring has been widely used to track pathogen dynamics (2), Illegal drug consumption (3), and to predict disease outbreaks (4). Wastewater is primarily treated in wastewater treatment plants (WWTPs), which employ various physical, chemical, and biological processes to remove organic matter and pollutants before releasing the effluent back into the environment.

Recent advances in shotgun metagenomics have enabled comprehensive profiling of wastewater microbial communities, including the characterization of the bacteriome (5) and virome (6), as well as the large-scale identification of antibiotic resistance genes (ARGs) (7, 8). Wastewater and WWTPs represent complex engineered ecosystems mixing microbial communities originating from households and diverse environmental sources. Furthermore, these environments include fluctuating nutrient concentration, aerobic and anaerobic niches, and seasonal variations, which collectively apply constant selective pressures that shape highly diverse microbial communities (9, 10). Hence, WWTPs are considered hotspots for ARGs and MGEs, facilitating horizontal gene transfer and the emergence of novel antibiotic resistance mechanisms (7, 11, 12). Additional studies have highlighted numerous biosynthetic gene clusters (BGCs) in activated sludge (AS) communities, most of which are novel, suggesting that wastewater microorganisms may represent an underappreciated reservoir of secondary metabolites with unknown functions (13, 14).

Generating protein families has emerged as the primary approach for grouping homologous protein sequences, exploring biodiversity, and inferring function across large datasets to annotate uncharacterized proteins, discover novel proteins, and predict microbial functions(15–19). Numerous generic databases provide curated protein annotations, structural information, and functional classifications, including UniProtKB (20), Protein Data Bank (PDB) (21), RefSeq (22), GenBank (23), IMG/M (24), Big Fantastic Database (25), MGnify (26), Pfam (27), InterPro (28), SMART (29), NMPFamsDB (30), metagRoot (31), MetaVR (32), UHGV (33), and COG (34). Databases more directly related to wastewater include the MiDAS 3 database, which provides taxonomic annotations of microbial communities in Danish WWTPs (35), and PharmaWaste, a literature-curated database of ARGs identified in WWTPs (36). Despite the growing number of metagenomic studies, to our knowledge, there is no resource focused on wastewater microbiomes at the protein family level.

Here we present WasteFams, the first database dedicated to the systematic exploration of protein families in wastewater metagenomic and metatranscriptomic studies at a global scale. WasteFams groups millions of proteins derived from wastewater datasets to provide novel insights into the functional landscape of wastewater microbial communities. By integrating scaffold-derived genomic context, we expanded the annotation of protein families, including their associations with MGEs, BGCs, and ARGs, offering a unique resource for the underlying mechanisms shaping wastewater microbial communities.

Notably, WasteFams is part of the EnvoFams portal (https://envofams.org), a broader platform dedicated to exploring and characterizing microbial and viral protein families across diverse ecosystems worldwide. The database can be found at https://envofams.org/wastefams or https://www.envofams.org/wastefams.

## MATERIAL AND METHODS

### Data Collection and protein family generation

Publicly available wastewater-related metagenomes, metatranscriptomes, and reference genomes were downloaded from the Integrated Microbial Genomes & Microbiomes (IMG/M) database (beginning 2024) (24). The compiled dataset included 580 assembled metagenomes, 132 metatranscriptomes, and 1,709 reference genomes collectively containing 105,798,843 proteins and 33,722,375 scaffolds. Samples were selected based on GOLD classification for Ecosystem Category (37), including “*Wastewater*”, “*Bioremediation*”, “*Sewage Treatment Plant*”, and “*WWTP*”. Regarding the Bioremediation-associated samples, only those that were related to wastewater were selected for further analysis.

To minimize redundancy, protein sequences were dereplicated using the MMseqs2 Linclust method with 100% sequence identity and coverage (38, 39). Four additional filtering steps were applied to retain only high-quality protein predictions. Specifically, *i)* scaffolds smaller than 500 bp were discarded to ensure that partial or truncated genes would be removed, *ii)* regions within 10 bp of scaffold boundaries were removed to diminish truncation events, *iii)* low complexity regions identified by tantan (40) were discarded, and *iv)* only protein sequences longer than 35 aa were retained. The dereplication and filtering steps resulted in 24,468,459 scaffolds, 1071 reference genomes, and 60,382,405 high-quality protein sequences.

For protein family generation, the MMseqs2 Linclust clustering algorithm was used in bidirectional mode with parameters set to 30% sequence identity and 80% bidirectional coverage between the query and the subject. Only protein families with 100 or more members were retained and aligned using MAFFT (41). To remove redundancy, the hhfilter (42) was applied to the aligned sequences with 95 % identity and 70% coverage. Finally, trimming was applied to minimize gaps with a custom script. Again, only protein families with 100 or more members were kept, and for those that passed this criterion, the filtered-out members were returned. The final dataset consists of 3,387 protein families from 633 datasets, 725,558 scaffolds, 1,151 isolates, and 1,068,945 protein sequences.

### Taxonomy

Scaffolded sequences associated with protein families were retrieved from the IMG/M database. Taxonomic classification was performed using Kraken2 (version 2.1.3) (43). To improve sensitivity and annotation depth (given the limitations of taxonomic algorithms like Kraken2, especially in handling eukaryotic sequences(44)), scaffolds that remained unclassified or were assigned to broad categories (e.g., ‘root’ or ‘cellular organisms’) were further analyzed using MMseqs2 taxonomy (version 13) (45). Last, to identify viral and plasmid signatures, any remaining unclassified scaffolds were processed through geNomad (version 1.8.1) (46).

### Biome and metadata collection

Habitat-related insights were obtained from environmental metadata using GOLD classification (37). Biome distributions for each protein family were inferred from sampling metadata associated with individual family members, based on the ‘Ecosystem Type’ field. Samples originated from a wide range of wastewater-related environments, including activated sludge, aerobic and anaerobic digesters, influent and effluent streams, mixed liquor, nutrient removal systems, sewage, sludge, industrial wastewater, terephthalate- and thiocyanate-associated systems, and unclassified wastewater sources.

### Structural prediction & annotation

Structural predictions were performed using AlphaFold3 (47) for protein families with at least 100 effective sequences after applying hhfilter to remove redundancy. For each family, five models were generated, and the one with the highest pLDDT score was selected. Only models classified as high (pTM ≥ 0.7) or medium-quality (pTM between 0.5 and 0.7) were retained.

To annotate the predicted structures, we performed Foldseek (48) against experimentally validated models in CATH (v4.4, December 2025) (49) and PDB (2024 release) (21), as well as experimental and predicted models in AlphaFoldDB (50). Significant matches were defined based on the following criteria: *i)* global TM ≥ 0.5, *ii)* for cases where the query was shorter than the target, query alignment score > 0.5, *iii)* for cases where the query was longer than the target, matches with target TM ≥ 0.5.

### Functional annotation

Protein families were annotated by searching the representative sequences against the Pfam database (v37, December 2024), which includes 21,979 families and 709 clans, using hmmsearch from HMMER 3.3.2 (51). Searches were performed using bit score thresholds of ≥25 at the sequence level and ≥22 at the domain level. For gene neighborhood analysis and annotation of all open reading frames (ORFs) within scaffolds, hmmsearch was performed with permissive thresholds, and domain hits were retained with e-values < 1e-5.

### Identification of CRISPR arrays and Cas systems

CRISPR–Cas systems were identified from the scaffold sequences associated with the protein families using CRISPRCasTyper v.1.8.0 (52) with the --prodigal meta option. For each scaffold, CRISPRCasTyper was used to detect CRISPR arrays and associated Cas genes, and to classify the corresponding CRISPR–Cas systems based on their internal reference models and scoring framework.

### ARG discovery

The scaffolds corresponding to protein families were used as input for protein-coding gene prediction using Prodigal (v2.6.3) (53). The resulting predicted proteins were screened for antibiotic resistance genes (ARGs) using the Resistance Gene Identifier (RGI, v6.0.4) with the Comprehensive Antibiotic Resistance Database (CARD) database (54). The most recent CARD reference data were downloaded in December 2025. ARG prediction was conducted using RGI in protein mode and DIAMOND (55) as the alignment tool.

To construct a high-confidence ARG dataset, filtering thresholds were defined based on the distribution of sequence-identity values for each resistance mechanism and model type, informed by biological knowledge of ARG families. Density distributions of % identity were examined to distinguish high-confidence hits from low-identity homologs. For protein homolog models (PHM), mechanism-specific thresholds were applied to balance sensitivity and specificity. Antibiotic inactivation (≥60%) and target protection (≥70%) showed clear, high-identity enrichment, allowing for moderate thresholds. In contrast, antibiotic target alteration required a stringent cutoff (≥95%) due to a large population of low-identity homologs, primarily driven by housekeeping variants of van-related genes, which would otherwise inflate false-positive assignments. Protein variant models (PVM) for target alteration were filtered at ≥70% identity to ensure reliable variant-level classification. Categories with limited and high-confidence distributions, such as antibiotic target replacement, were retained without additional filtering. Antibiotic efflux predictions derived from PHM were excluded, as previously reported(56), as their identity distributions were broad and could not be reliably distinguished by conserved housekeeping transporters. In contrast, overexpression models (OEM), including efflux-associated entries, were retained as they represent experimentally supported resistance mechanisms.

### MGE annotation

The predicted proteins were additionally screened for MGEs using a curated MGE database (57). Coding sequences from the MGE database were predicted using Prodigal (53), and the resulting proteins were used to construct a custom reference database with DIAMOND. Predicted protein sequences from the metagenomic assemblies were queried against this MGE protein database using DIAMOND BLASTP (55). Searches were performed with an e-value threshold of 1 × 10^−5^. Output files included percent identity, alignment length, query length, subject length, e-value, and bitscore. Alignments were filtered to retain only hits with at least 30% amino acid identity and at least 80% query coverage. For each ORF, only the best-scoring hit based on bitscore was retained to avoid redundant assignments.

### Gene neighborhood analysis

For the case study, bedtools window was used to examine a ±1000 bp genomic window for all protein family members. Neighboring ORFs were represented based on their best Pfam prediction. Only neighbors that occurred more than 5 times next to a protein family gene were kept and visualized into a network using Cytoscape (58). Graphical representation of selected scaffolds was performed with the Bioconductor package ggggenes v.0.6.0.

### BGC identification

BGCs were identified on scaffolds associated with protein families and longer than 5,000 bp, using antiSMASH v8 (59) in bacterial mode. Gene prediction was performed through Prodigal. The analysis included ClusterHMMER, active site finder, known-cluster and subcluster comparisons, MIBiG comparison, TIGRFAM annotation, Pfam2GO mapping, RRE detection, and TFBS prediction.

Using genomic coordinates, we mapped protein family members to BGCs that encode them. Proteins encoded by genes located within BGC core regions were classified as core members, whereas those encoded in flanking regions were classified as accessory members. Our case study focused exclusively on core BGC members predicted to produce RiPPs and identified those that repeatedly co-occur within the same clusters. However, the WasteFams database provides functionality for exploring both core and accessory members across all product classes.

### Database implementation and structure

The WasteFams platform is built using the ASP.NET Core Model-View-Controller (MVC) framework, with DuckDB (60) serving as its database management system. The backend is implemented in C#, while the frontend is developed using HTML, CSS, and JavaScript.

The system is organized into components, including Models, Views, Controllers, Services, and Factories. Models, defined as C# classes, represent the underlying data structures. Views utilize Razor (.cshtml) templates combined with C# logic to dynamically generate HTML content. Controllers act as intermediaries, processing user requests and coordinating interactions between Models and Views. Services handle data processing, while Factories contain the application logic.

WasteFams integrates a comprehensive selection of sequence analysis tools. Protein sequence searches are performed using DIAMOND (61), while HMM-based searches are also supported through HMMER (62). Additional functionality includes pattern-based sequence searches and structural similarity searches using Reseek (63) for 3D structure models. All computational tasks are managed by a workload manager, in which each query is queued, assigned a unique job identifier, and can be revisited by users to retrieve results.

The platform also incorporates several visualization and exploration tools. These include a Feature Viewer (64) for annotated sequence features, MSAViewer (65) for multiple sequence alignment visualization, the Mol* viewer (66) for interactive 3D molecular structure rendering, and a genome browser powered by CGView (67) for genomic data visualization. Additionally, OpenStreetMap is integrated to support geospatial data visualization.

## RESULTS

### Database components

WasteFams is a repository cataloging 3,887 protein families that contain ≥100 sequences derived from 580 metagenomic and 132 metatranscriptomic samples, as well as 1,709 reference genomes associated with wastewater and WWTP ecosystems. In total, ∼106 million predicted proteins, and ∼33 million scaffolds were processed to construct non-redundant protein families that were subsequently characterized using functional and structural annotations. For each family, MSAs and HMMs were generated, and structural models were predicted with Alphafold3 (47). Accordingly, taxonomic classifications and associated ecosystem metadata were included. The resulting protein families were further linked to ARGs, MGEs, and BGCs, allowing exploration of key functional traits associated with wastewater microbiomes (Figure 1).

**Figure 1:**
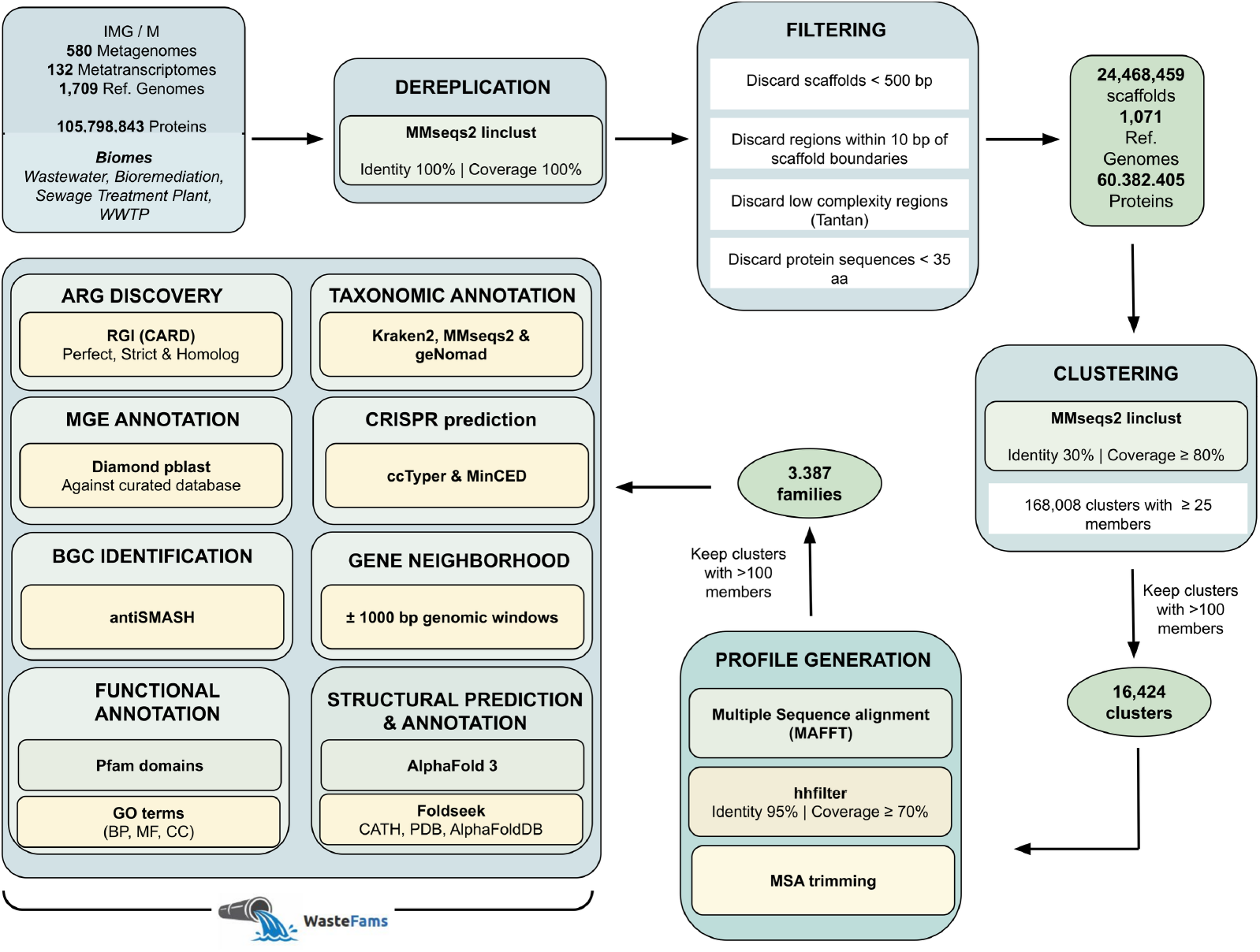
Data preprocessing and protein family construction pipeline. The workflow includes dereplication, sequence filtering, clustering, and profile generation to construct protein families from wastewater-related metagenomic, metatranscriptomic, and reference datasets. For each family, multiple sequence alignments and profile HMMs were generated, and structural models were predicted using AlphaFold3. Protein families were further annotated using functional, taxonomic, and genomic context analyses, including the detection of ARGs, MGEs, and BGCs.

### The WasteFams interface

The WasteFams platform is organized around a navigation bar that provides access to sections including Browse, Sequence Search Tools, Statistics, Downloads, and Help pages, enabling seamless navigation to all database features (Figure 2A). The main page also displays key statistics and summary metrics to provide an overview of WasteFams’ characteristics.

**Figure 2:**
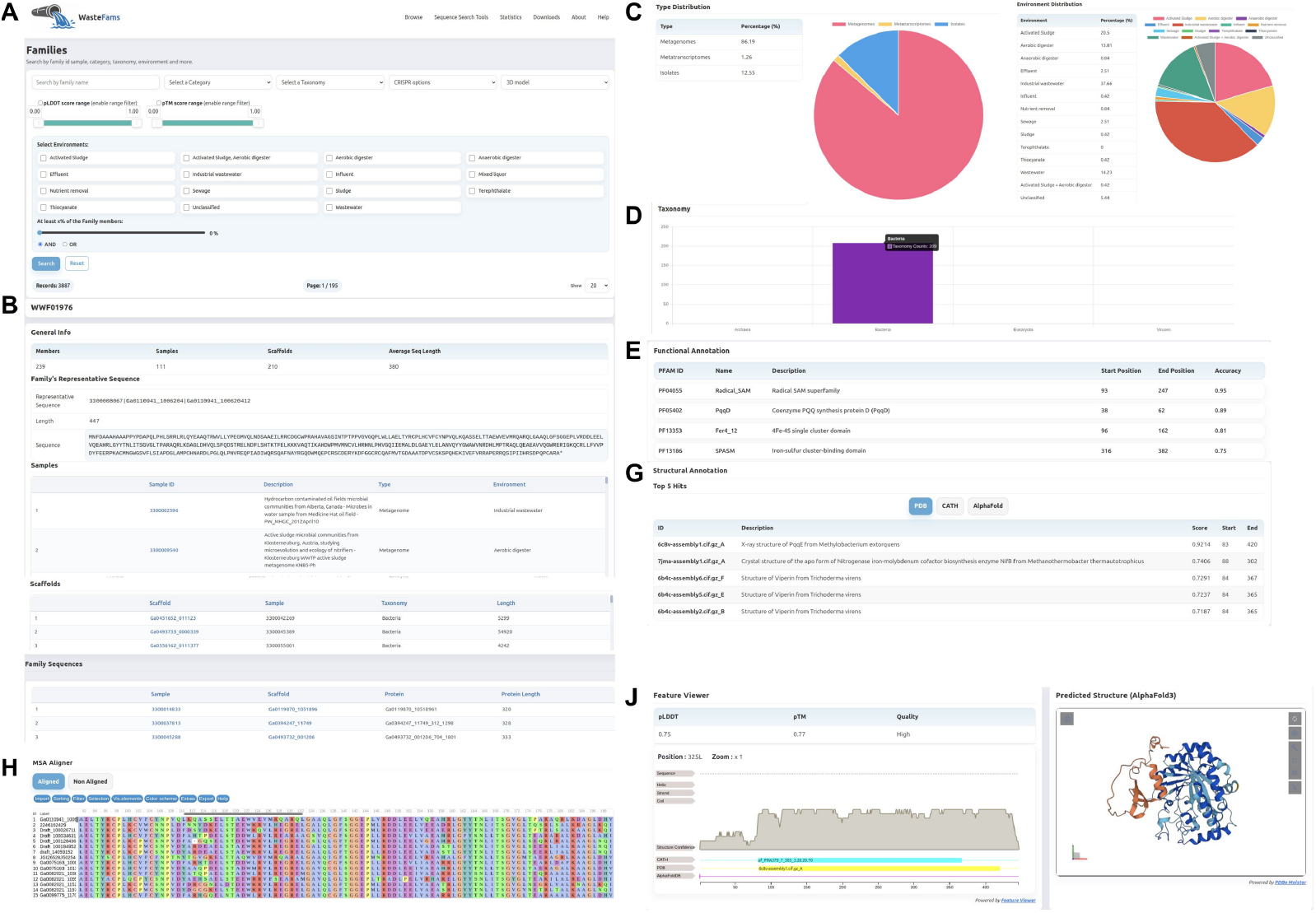
WasteFams protein family interface. The Browse Families page offers free-text search and filtering by sequencing category, taxonomy, environmental distribution, structural prediction metrics, and family size. Upon selection, each protein family is displayed with its identifier, number of members, associated datasets and scaffolds, average sequence length, and representative amino acid sequence. Additional tables report Pfam functional domains and top structural matches. Interactive viewers include an MSA with customization and FASTA export; a sequence logo from the profile HMM with export options; a linear feature map of topology, secondary structure, conservation, and domain annotations; and an embedded AlphaFold3 3D model. Interactive graphical summaries include pie charts showing the distribution of sequencing categories and environmental origins of family members, as well as a bar chart illustrating their taxonomic distribution. All alignments, profiles, structural models, and annotation tables are available for download in standard bioinformatics formats.

### Browse families: data, visualization, and metadata context

The Browse → Families section provides a comprehensive query interface supporting both simple and advanced searches. Protein families can be queried using a free-text search by family identifier, while additional filters are available through dropdown menus, including sequencing category (metagenome-only, metatranscriptome-only, mixed or all families), taxonomy, association with CRISPR–Cas systems, and 3D model prediction quality. Range sliders enable filtering based on pLDDT and pTM scores derived from AlphaFold predictions. Users can focus on selected environments using checkboxes and define the desired proportion of family members present in those habitats. Results are displayed in an interactive table summarizing key attributes of each protein family, including the number of members, associated datasets and scaffolds, and structural model quality. The table supports sorting across all columns, facilitating efficient exploration of the protein families (*Figure 2A*).

Upon selecting a protein family, a dedicated page displays its unique identifier along with summary statistics, including the total number of member sequences, associated datasets and scaffolds, and the average sequence length. The representative sequence is also provided, including its identifier, protein length, and amino acid sequence. All identifiers are linked to IMG/M. Two tables list the associated datasets and scaffolds, coupled with their respective metadata (*Figure 2B*). Moreover, graphical summaries are provided, including two pie charts showing the proportion of sequencing approaches associated with each family member and the distribution of protein families across different habitats (*Figure 2C*). A bar chart illustrating the distribution of protein family members across major microbial groups (bacteria, archaea, viruses, and protozoa) is also included (*Figure 2D*).

Functional annotation of each protein family is provided through Pfam predictions, including domain positions and scores (*Figure 2E*). Structural annotation reports the top five PDB, CATH, or AlphaFoldDB matches for the family’s representative sequence, including TM-scores and alignment coverage to indicate structural similarity (*Figure 2G*). In addition, a table lists protein family members associated with known MGEs (where available), along with the matching domains and corresponding alignment statistics.

Next, several interactive tools are provided to explore protein family member sequences. MSAs can be viewed in either full mode, showing all sequences in the alignment, or seed mode, showing only the representative core sequences used to define the alignment, with options for custom coloring, identity and occupancy thresholds, and motif or regular-expression searches. (*Figure 2H*). An adjacent sequence logo viewer renders the family’s profile HMM as an interactive plot of residue probabilities (each position being clickable to reveal detailed scores). Both the MSA sequences and the underlying HMM can be exported in FASTA, HMMER, or HH-suite format accordingly. Per-residue annotations, including transmembrane topology, secondary structure elements, conservation profiles, and domain predictions (e.g., Pfam), are mapped onto the representative sequence using a linear feature viewer (*Figure 2J*). When available, an AlphaFold model is provided as an interactive, rotatable three-dimensional ribbon representation of the predicted structure. Structural information can be downloaded in CIF format (*Figure 2J*). Finally, an interactive map illustrating the locations of the corresponding sampling sites (when coordinates are available in IMG) is provided.

### Browse scaffolds and search tools

The Browse → Scaffolds section provides a comprehensive query interface that allows users to navigate through samples and their associated scaffolds. Dropdown menus enable filtering of scaffolds by specific environments and microbial groups, as well as by the presence or absence of CRISPR arrays. A sortable table that lists each scaffold with its length, number of family members, taxonomy, and associated habitat is provided.

At the top of the scaffold view, a summary table provides general information for each scaffold, including sample ID, scaffold ID, sample name, sequence length, biome, and taxonomic assignment. Additional features are provided to further characterize the genomic context and its association with the protein families (*Figure 3A*).

**Figure 3:**
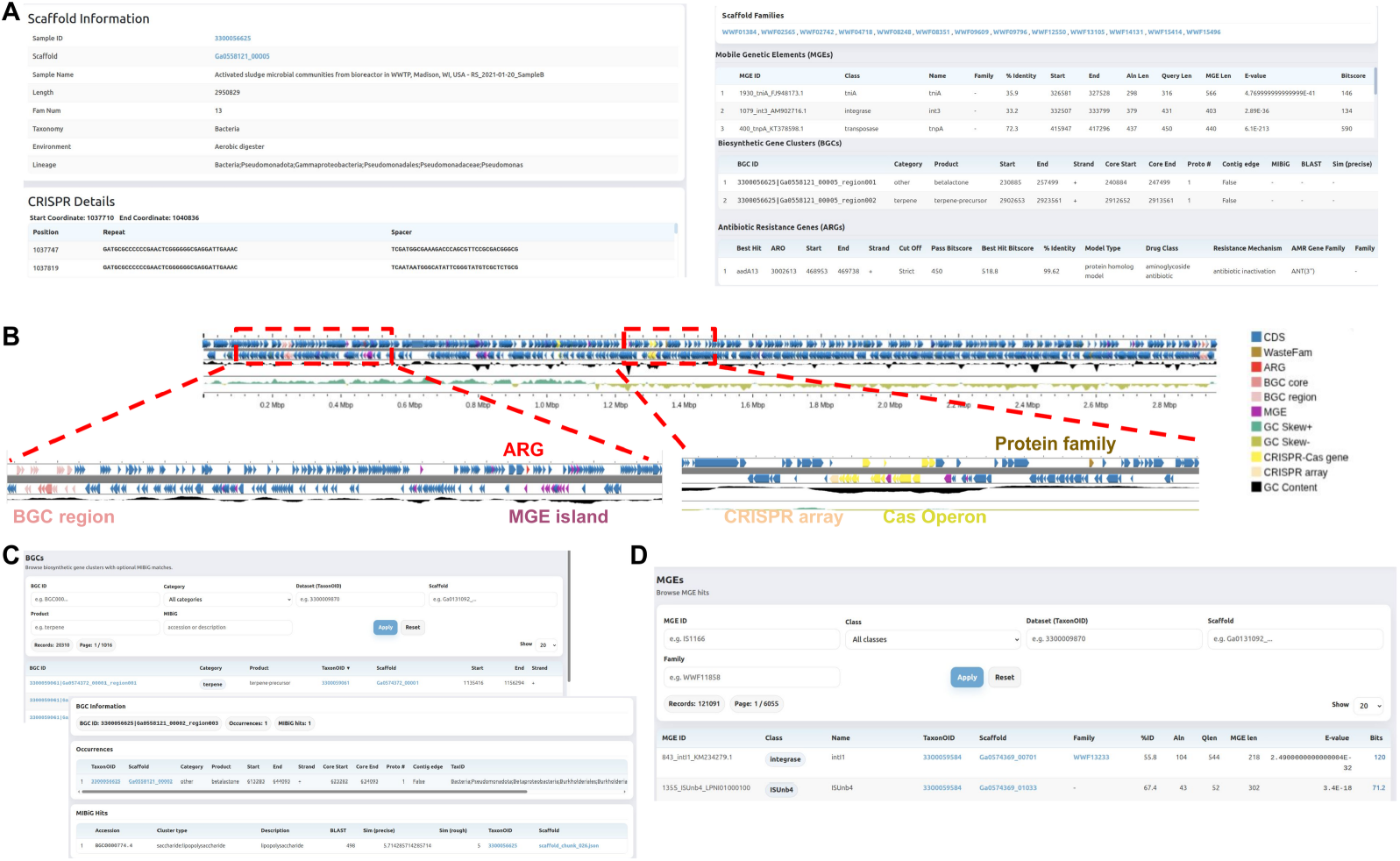
WasteFams interface and browse tabs. (A) The Browse Scaffolds page leads to a page containing a table with general information, including sequence length, the number of associated protein families, environmental metadata, and taxonomic assignment. Additional tables report CRISPR arrays, identified protein families, the MGE annotations, and BGCs. (B) A scaffold viewer provides the genomic context of the scaffold, containing distinct colored labels for protein families, CRISPR arrays & Cas operons, MGEs, BGCs, ARGs, and coding DNA sequences (CDS). (C) The BGC module supports free-text search and filtering by BGC class, product type, datasets, scaffolds, and similarity to known clusters in MIBiG. Each BGC entry links to detailed pages including antiSMASH predictions and MIBiG annotations. (D) The MGE module enables free-text search and filtering by dataset, scaffold, MGE class, and associated protein families. Results are presented in a table that summarizes MGE annotations and DIAMOND-derived sequence similarity metrics.

CRISPR arrays are reported in a dedicated table, including the positions of the corresponding repeats and spacers. A distinct section lists the protein families that are identified in the specific scaffold. Next, the ORFs matching known MGEs are listed along with alignment statistics, including alignment length, E-values, and bit scores. Identified BGCs are summarized in a dedicated table that reports cluster category, predicted product type, and genomic coordinates of core regions, along with similarity to known clusters in MIBiG. In addition, predicted ARGs are presented in a separate table that includes associated metadata, such as bit score, model type, drug class, resistance mechanism, and AMR gene family. Finally, an integrated genome viewer summarizes all annotated genomic features (CRISPR arrays/operons, MGEs, ARGs, BGCs), enabling seamless navigation across the scaffolds (*Figure 3B*).

Additional sections of WasteFams provide a broader overview of reference genomes and curated MGE and BGC datasets. The MGE section allows users to explore specific MGE classes and monitor MGE presence in metagenomic samples(*Figure 3C*). The BGC section provides a comprehensive list of identified clusters, including their category, product class, and associated scaffolds and samples. Each BGC entry links to detailed information, including similarity to experimentally validated clusters from the MIBiG database (*Figure 3D*).

### Sequence search and visualization tools

To expand the usability of WasteFams, a suite of tools is provided to query user-submitted sequences against the entire database. Sequence similarity searches are performed using DIAMOND for rapid alignment-based comparisons, while HMM-based searches are performed with HMMER to identify profile matches. Both tools allow users to customize key parameters, including score cutoffs, substitution matrices, and gap penalties. A dedicated Pattern Search interface supports PROSITE-style motif queries or arbitrary regular expressions, enabling the detection of conserved sequence features across protein families. In addition, structural similarity searches are implemented using Reseek (63), with adjustable sensitivity modes, E-value thresholds, and query coverage. All result tables can be readily downloaded for downstream analysis.

Conclusively, WasteFams is a user-friendly platform for exploring 3,887 protein families across hundreds of metagenomic and metatranscriptomic datasets from wastewater-related habitats. By linking the protein families to MGEs, BGCs, ARGs, and CRISPR elements, the platform provides an enriched functional context to support in-depth analysis. Furthermore, the integration of advanced search tools (HMMER, DIAMOND, Reseek, and PROSITE/regex), along with interactive visualization and customizable data export options, enables hypothesis generation and systematic exploration of wastewater-associated microbial functions.

### Data contents and case studies

General properties of the identified protein families were examined (*Supplementary Figure 1A-D*). Most proteins ranged between approximately 100 and 300 amino acids, while the protein length distribution was right-skewed overall (*Supplementary Figure 1A)*. Consistent with this observation, the average protein length per family was primarily distributed between ∼200 and 1,400 amino acids, with a peak around 300–400 aa (*Supplementary Figure 1B)*. Protein family sizes were skewed toward smaller groups, with most families containing approximately 300–400 members, but we also observed families with 1,556 and 1,823 members (*Supplementary Figure 1C)*. Metagenomic and metatranscriptomic scaffolds typically ranged from ∼2,500 to 5,000 bp, with several thousand exceeding 500,000 bp.

Taxonomic annotation of the scaffolds associated with the identified protein families revealed that approximately 99% of the sequences were assigned to Bacteria (*Supplementary Figure 1E)*. The microbial community was dominated by members of the phylum Pseudomonadota (Proteobacteria), including families such as Pseudomonadaceae, Lysobacteraceae, Enterobacteriaceae, Nitrobacteraceae, and Rhizobiaceae. Members of the phylum Bacteroidota were also abundant, including representatives of the class Bacteroidia and genera such as Flavobacterium. Our observations demonstrate communities that represent both human activity and the biological processes that occur in WWTPs and are consistent with previous studies reporting that Proteobacteria and Bacteroidota dominate microbial communities in sewage and wastewater treatment plants (5, 68).

Next, we examined the presence of ARGs using the RGI tool against the CARD database (54). We found that the microbial communities in wastewater were characterized by ARGs conferring resistance to multiple antibiotic classes, as has been previously reported (7, 56). Notably, Pseudomonadota, Bacillota, Actinomycetota, and Bacteroidota were the most frequently ARG-associated phyla (*Supplementary Figure 1F)*.

These scaffolds were further examined for the presence of CRISPR arrays and associated Cas operons. In total, 3,368 CRISPR arrays were identified across 3,367 scaffolds, together with 2,195 Cas operons. Integration of these annotations with protein family assignments revealed that 15 protein families contained members encoded within Cas operons. The functional association of these families with CRISPR–Cas systems was supported by Pfam domain annotations consistent with Cas-related proteins.

The biome distribution of the protein families, based on the GOLD ecosystem classification, revealed strong representation in industrial wastewater, activated sludge, wastewater, and aerobic digester environments (*Supplementary Figure 1G*). This widespread distribution suggests that many of the identified proteins play essential roles in core components of wastewater microbial ecosystems, but it also reflects the continuous mixing and recycling of microbial communities throughout the wastewater treatment process.

To further characterize the identified protein families, we predicted the structures of the representative sequences using Alphafold3. The majority of the predicted models were of high confidence (pTM > 0.7) or medium confidence (pTM > 0.5), indicating that most proteins adopt well-defined structural folds (*Supplementary Figure 1H*). Collectively, our analyses highlight the broad taxonomic, functional, ecological, and structural diversity of wastewater metagenomes, offering multiple avenues for exploration for WasteFams users.

### Case study I: Characterization of MGEs associated with protein families

Multiple studies have highlighted that Wastewater microbiomes are rich in MGEs that facilitate the horizontal transfer of genomic features and ARGs (7, 56, 69). To investigate the contribution of the WasteFams protein families to these mechanisms, we searched for MGE-associated proteins using DIAMOND BLAST against a curated MGE database (57). In total, we identified 121,091 MGEs across 62,433 scaffolds, with 69 families containing 10,541 protein members matching known MGEs. The most abundant class was transposases, followed by integrases and IS91-associated proteins (*Figure 4A*), consistent with previous studies (56, 70). Even though transposition was dominant, we identified several classes of insertion sequences, as well as recombination proteins and plasmids at lower frequencies, suggesting diverse mechanisms of genomic mobility across wastewater microbial communities.

**Figure 4:**
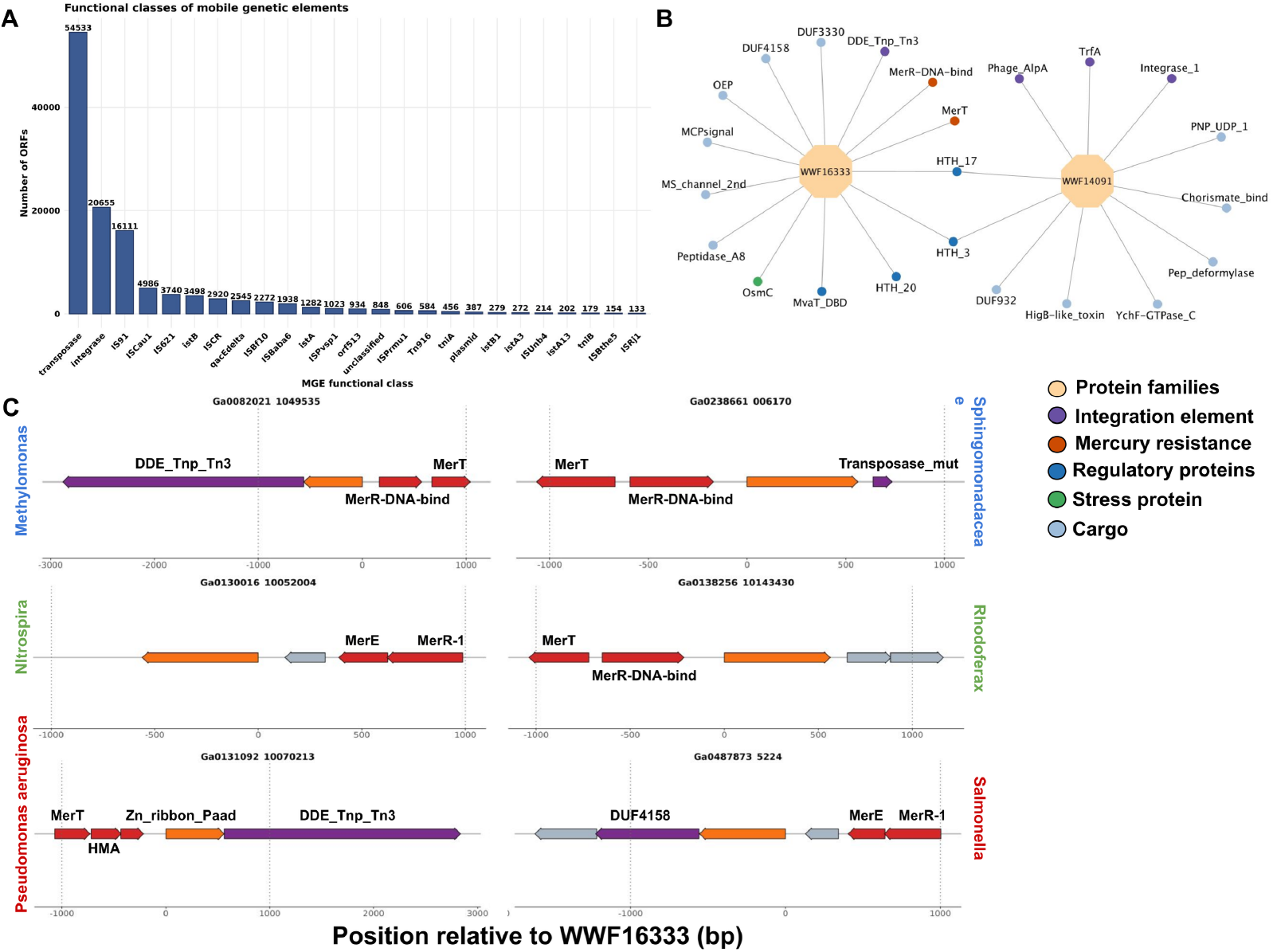
Characterization of MGEs associated with WasteFams protein families. (A) Bar plot showing the distribution of functional classes of MGE-associated proteins identified using DIAMOND BLAST against a curated MGE database. (B) Network representation of selected protein families and their associated functional domains identified within genomic neighborhoods. Nodes represent protein families or functional domains, while edges indicate co-occurrence within ±1,000 bp genomic windows. Only domains that were detected more than 5 times were visualized in the network. Domain annotations include integrative elements (purple), mercury resistance domains (red), regulatory proteins (blue), stress-response proteins (green), and cargo genes (light blue). (C) Graphical representation of selected genomic scaffolds illustrating the co-localization of protein family WWF16333 with mercury resistance genes and integration elements across different bacterial taxa, in a ±1000 bp genomic window.

To further decipher the genomic context of these elements, we examined the gene neighborhoods within ±1,000 bp of the corresponding protein family members and performed network analysis of associated protein domains. We discovered recurrent co-occurrence patterns (>5 occurrences) among several protein families and domains associated with integrative elements, regulatory proteins, and metabolic cargo genes. Among these, the protein families WWF16333 and WWF14091 were repeatedly detected on the same scaffolds, along with integration elements, metabolic cargo genes, and regulatory proteins (*Figure 4B*). The protein family WWF16333 consists of 374 conserved members containing resolvase and HTH7 domain, while WWF14091 includes 227 members with DNA-binding domains and phage integrase-related regions. Notably, both families share common protein interactants with regulatory domains, including HTH_7 and HTH_3, which are commonly associated with MGEs (*Figure 4B*). These observations suggest that WWF16333 and WWF14091 may cooperate in multiple horizontal transfer events within wastewater microbiomes.

Moreover, we observed that members of the WWF16333 family are frequently found in close proximity to mercury resistance genes (*Figure 4B*). Mercury resistance is a common adaptive trait in wastewater environments, leading to the emergence of resistant bacteria that tolerate and degrade heavy metals (71). Visual inspection of selected scaffolds containing mercury-resistance genes revealed colocalization of WWF16333 genes with transposases and integrative elements, suggesting a potential role for this protein family in the horizontal gene transfer of mercury-resistance genes. This mobilome–resistance axis was detected across diverse taxa, including environmental bacteria (Methylomonas, Sphingomonadaceae), wastewater-associated degraders (Nitrospira, Rhodoferax), and clinically relevant pathogens (Pseudomonas aeruginosa, Salmonella). Collectively, our results demonstrate that information on the WasteFams protein family, along with the corresponding genomic context, can provide novel insights into horizontal gene transfer mechanisms within wastewater microbiomes.

### Case study II: Functional annotation of protein families that contribute to BGCs

Studying environmental BGCs provides insights into the discovery of natural secondary metabolites with potential biotechnological applications as antibiotics, antifungals, or anticancer agents. Using antiSMASH, we predicted 20,311 BGCs, of which 17,157 originated from metagenomic scaffolds and 129 from metatranscriptomic scaffolds associated with protein families. Among the detected clusters, 42% corresponded to terpene BGCs, 22% to ribosomally synthesized and post-translationally modified peptides (RiPPs), 12% to polyketide synthases (PKS), and 8% to non-ribosomal peptide synthetases (NRPS), while the remaining 15% belonged to other classes (*Figure 5A*).

**Figure 5:**
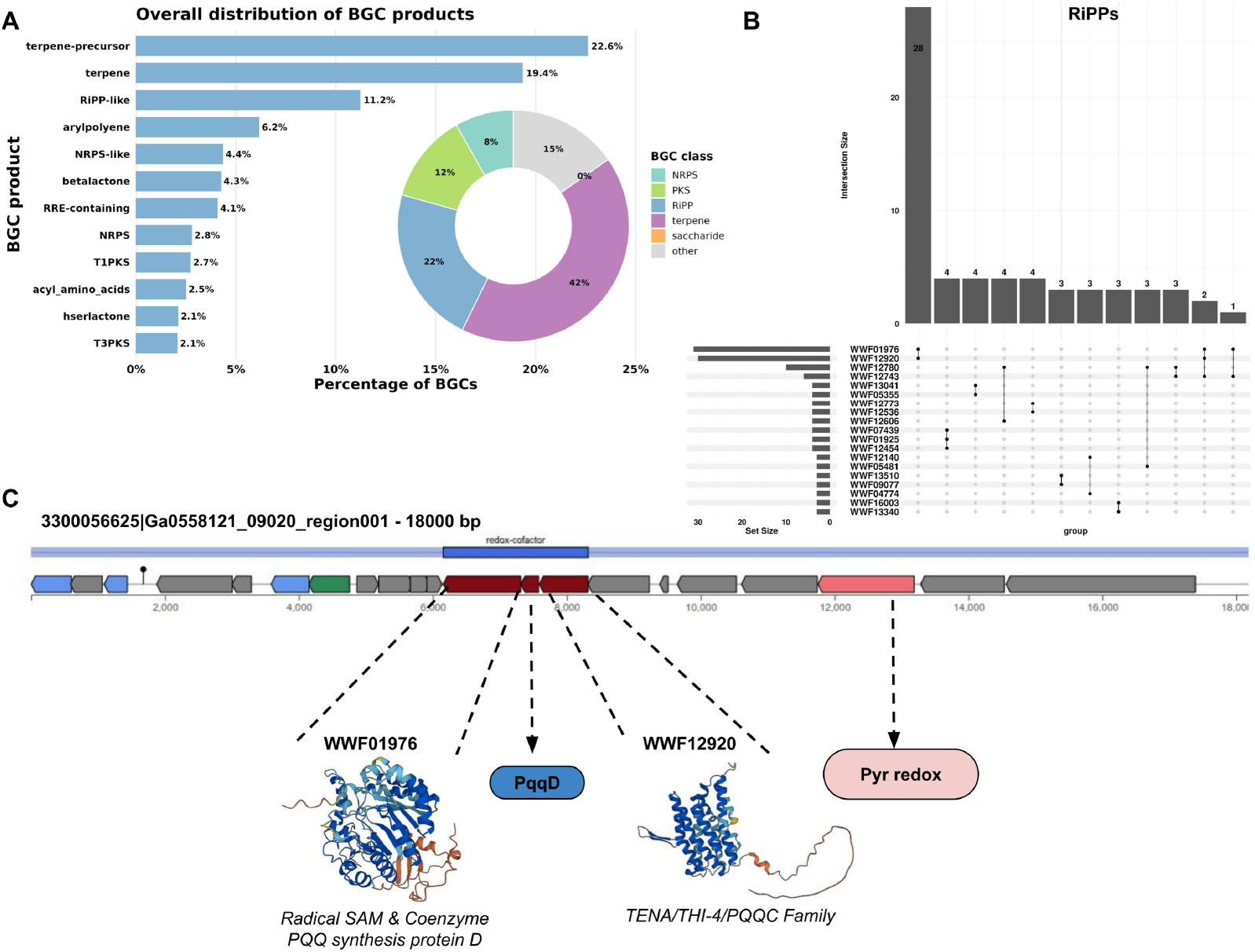
Identification and characterization of BGCs associated with WasteFams protein families. (A) Bar chart showing the distribution of the most frequently predicted BGC product types identified using antiSMASH, while the pie chart summarizes the relative distribution of major BGC classes. (B) UpSet plot illustrating the co-occurrence of protein families consistently found within the same RiPP cluster cores. (C) Genomic organization of a representative BGC identified in scaffold 3300056625|Ga0558121_09020 (33,384 bp). The predicted BGC spans approximately 18 kb and contains genes encoding members of the protein families WWF01976 and WWF12920, as well as proteins associated with pyrroloquinoline quinone (PQQ) biosynthesis. Structural predictions for representative proteins from these families are shown below the cluster, highlighting domains associated with PqqD and PQQ synthase.

RiPP BGCs typically include genes encoding a precursor peptide, along with enzymes responsible for its post-translational modifications. and in some cases, leader peptidases and transporters. The precursor peptide is synthesized by ribosomes and subsequently modified to yield the mature compound (72). RiPPs represent an important class of natural products with significant antibacterial activities and great potential for the discovery of novel antibiotics (73).

To explore the functional context of the predicted BGCs, we investigated protein families that repeatedly co-occurred within RiPP clusters. Notably, we found that proteins belonging to the families WWF01976 and WWF12920 were repeatedly detected within the same RiPP BGCs, suggesting that these families may represent conserved components of a shared biosynthetic pathway in wastewater (*Figure 5B*). The family WWF12920 consists of 381 members, most of which are found in industrial wastewater samples (43.83%), and contains a domain related to Pyrroloquinoline quinone synthase, based on foldseek annotations and Pfam hits. Accordingly, the WWF01976 family comprises 239 members, predominantly detected in industrial wastewater (37.66%), and contains domains associated with Coenzyme PQQ synthesis protein D (PqqD) and the radical SAM superfamily. These proteins are core components of BGCs involved in the biosynthesis of Pyrroloquinoline Quinone (PQQ), a redox-active quinone known to protect against oxidative stress and proposed for therapeutic applications in metabolic disorders and cancer (74).

A closer inspection of the scaffold 3300056625|Ga0558121_09020 (33,384 bp) revealed a novel BGC (no strong matches against MiBiG database) of approximately 18,000 bp. The cluster contains a redox-cofactor biosynthetic core that encodes members of WWF01976 and WWF12920, as well as a PqqD protein. A gene encoding Pyr redox is located adjacent to the core BGC region and may play a supporting role in redox metabolism (*Figure 5C*). Collectively, our insights suggest that this locus represents a novel PQQ-related biosynthetic pathway with conserved components.

Together, these observations demonstrate how navigation through WasteFams enables the identification and functional characterization of protein families associated with previously uncharacterized BGCs, paving the way for more efficient exploration of novel secondary metabolite pathways in wastewater metagenomes.

## DISCUSSION

By integrating global metagenomic, metatranscriptomic, and reference-genome data from wastewater environments, WasteFams provides the first unified and annotated resource for exploring protein-family diversity in wastewater microbiomes. Our catalog of 3,887 protein families is enriched with structural, functional, and ecological annotations and further expanded with information for their association with MGEs, BGCs, and ARGs. Through a user-friendly interface, WasteFams enables researchers to seamlessly explore the functional landscape of wastewater microbiomes and examine their evolutionary dynamics and the development of resistance mechanisms. Given the complexity of wastewater environments, the database facilitates the identification of protein families associated with human pathogens, environmental degraders, and microbial communities involved in WWTP processes. With the increasing number of global initiatives leveraging wastewater surveillance, including the European Urban Wastewater Treatment Directive, WasteFams provides a valuable platform for studying microbial function in wastewater ecosystems. Together, these features position WasteFams as a valuable resource for exploring wastewater microbiomes and for uncovering novel microbial functions, resistance mechanisms, and secondary metabolite biosynthetic pathways.

## ACKNOWLEDGEMENTS

This work was supported by computational resources provided by the ELIXIR-GR HYPATIA Cloud infrastructure.

## CONFLICT OF INTEREST

None declared.

## DATA AVAILABILITY

All data generated within WasteFams is fully accessible and downloadable. Multiple sequence alignments and protein family profiles can be exported in standard FASTA and HMMER or HH-suite formats, respectively. Three-dimensional structural models are available in CIF format, while annotation and metadata tables can be downloaded as TSV files. WasteFams is part of the EnvoFams portal (https://envofams.org), a broader platform dedicated to exploring and characterizing microbial and viral protein families across diverse ecosystems worldwide. The database is available at https://envofams.org/wastefams or https://www.envofams.org/wastefams.

## FUNDING

A.G, I.C, E.A, and G.A.P. were supported by the Hellenic Foundation for Research and Innovation (H.F.R.I.) under the “Third Call for H.F.R.I. Research Projects to support faculty members and researchers” [23592-EMISSION]; F.A.B. was supported by the Hellenic Foundation for Research and Innovation (H.F.R.I.) under the “4th Call for H.F.R.I. Research Project to support Postdoctoral Researchers” [28787-VIROMINE]; I.G.-S. was supported by startup funds from the Penn State College of Medicine and the University of Texas at Austin. The work conducted by N.C.K. in the US Department of Energy Joint Genome Institute (https://ror.org/04xm1d337) was supported by the US Department of Energy Office of Science user facilities, operated under contract number DE-AC02-05CH11231.

## Notes

https://envofams.org/wastefams

### Competing Interest Statement

The authors have declared no competing interest.

