## Supplementary File for "WasteFams: A database of protein families from global wastewater microbiomes"

### SUPPLEMENTARY DATA

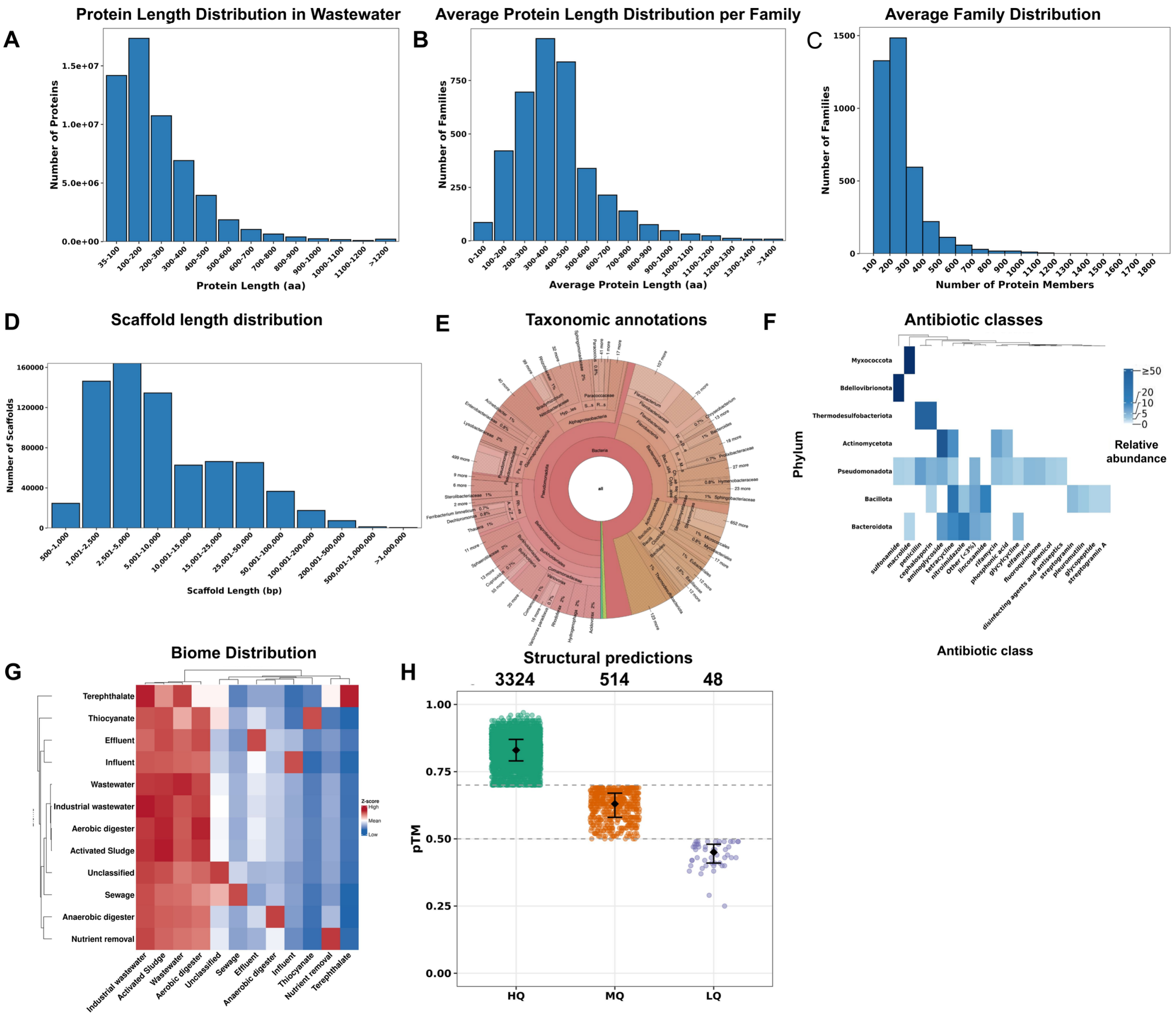


**Supplementary Figure 1: Distribution and characteristics of wastewater protein families.** (A-D) Histograms showing the distribution of (A) predicted protein lengths, (B) average protein length per family, (C) protein family sizes, and (D) scaffold lengths derived from metagenomic and metatranscriptomic assemblies. (E) KRONA visualization showing the taxonomic annotation of scaffolds associated with the protein families. (F) Heatmap demonstrating the relative abundance of detected ARGs across the most frequent ARG-associated phyla. Each cell represents the percentage of ARG hits within a phylum assigned to a given antibiotic class. Low-abundance classes (<3% per phylum) were collapsed into "Other (<3%)". (G) Heatmap showing the distribution of protein families across wastewater ecosystem types based on GOLD classification. Values are represented as z-scores to normalize differences in protein family abundance across biomes. (H) Scatter plot showing the pTM scores of structural predictions for representative protein family sequences using Alphafold3. Predicted structures were categorized as high quality (HQ, pTM > 0.7), medium quality (MQ, pTM > 0.5), and low quality (LQ, pTM ≤ 0.5).
